# DNA-based copy number analysis confirms genomic evolution of PDX models

**DOI:** 10.1101/2021.01.15.426865

**Authors:** Anna C. H. Hoge, Michal Getz, Rameen Beroukhim, Todd R. Golub, Gavin Ha, Uri Ben-David

**Affiliations:** Public Health Sciences Division, Fred Hutchinson Cancer Research Center, Seattle, WA, USA; Department of Human Molecular Genetics and Biochemistry, Faculty of Medicine, Tel Aviv University, Tel Aviv, Israel; Broad Institute of Harvard and MIT, Cambridge, Massachusetts, USA; Dana-Farber Cancer Institute, Boston, Massachusetts, USA; Harvard Medical School, Boston, Massachusetts, USA

## Abstract

We previously reported the genomic evolution of the copy number (CN) landscapes of patient-derived xenografts (PDXs) during their engraftment and passaging^1^. Woo et al. argue that the CN profiles of PDXs are highly conserved, and that the main conclusions of our paper are invalid due to our use of expression-based CN profiles^2^. Here, we reassess genomic evolution of PDXs using the DNA-based CN profiles reported by Woo et al. We find that the degree of genomic evolution in the DNA-based dataset of Woo et al. is similar to that which we had previously reported. While the overall Pearson’s correlation of CN profiles between primary tumors (PTs) and their derived PDXs is high (as reported in our original paper as well), a median of ~10% of the genome is differentially altered between PTs and PDXs across cohorts (range, 0% to 73% across all models). In 24% of the matched PT-PDX samples, over a quarter of the genome is differentially affected by CN alterations. Moreover, in matched analyses of PTs and their derived PDXs at multiple passages, later-passage PDXs are significantly less similar to their parental PTs than earlier-passage PDXs, indicative of genomic divergence. We conclude that genomic evolution of PDX models during model generation and propagation should not be dismissed, and that the phenotypic consequences of this evolution ought to be assessed in order to optimize the application of these valuable cancer models.

## Main text

Woo et al. assembled CN profiles of 1,451 samples corresponding to 509 PDX models^2^. CN profiles were evaluated using several methods, including SNP arrays, WES, WGS and RNAseq. As expected, RNA-based CN inference was found to be less accurate than that based on DNA measurements. To assess the similarity between PTs, early-passage PDXs and later-passage PDXs, Woo et al. relied mostly on a Pearson’s correlation analysis of the log_2_(CN ratio) values across the genome. They report high correlations between the CN profiles of PTs and PDXs (median correlation PT-PDX = 0.950), and between matched pairs of PDXs (median correlation PDX-PDX = 0.964). Furthermore, the authors report that the CN profiles remained highly stable throughout *in vivo* passaging, so that high-passage PDX models represented the CN landscapes of the PTs as well as early-passage PDXs. Based on these results, the authors suggest that our previous results were an artifact of inaccurate expression-based CN profiling.

The overall correlation of CN landscapes between PTs and PDXs and between related PDX samples is not surprising, and is consistent with our previous findings^1^. However, at the individual tumor level, we previously found that a median of 12.3% of the genome was differentially altered within four passages of PDX models (range, 0% to 59%). Consistent with our previous study, several recent papers have also reported genomic evolution throughout the engraftment and passaging of PDXs^3–10^. Intrigued by the apparent discrepancies between these previous results and those reported by Woo et al., we re-analyzed the CN data presented in their study.

We focused only on DNA-based (SNP arrays, WES, and WGS) CN profiles to avoid any potential issues with expression-based CN inference. In Woo et al.’s dataset, this included 33 PDX cohorts (representing 16 cancer types) with at least one pair of matched samples that had high-quality DNA-based CN calls^2^. We divided the genome into bins of 1 Mb length, and evaluated the discordance of each bin between PTs and PDXs, using conservative thresholds for ‘discordance’ (**Methods**). As an alternative approach, we predicted integer CN using ichorCNA^11^, which explicitly accounted for tumor impurity in the primary tumors and potential mouse contamination in the PDX samples, and focused downstream analysis on clonal events to allow for a conservative discordance comparison (**Methods**). Using both approaches, the fraction of the genome discordant between two samples was defined for each PDX model. Woo et al. reported that most discordant CNAs corresponded to small focal events (average size: 1.53 Mb)^2^, in contrast to our previous findings that CN evolution often involved entire chromosome arms and whole chromosomes. We therefore also evaluated the number of discordant chromosome-arm CNAs in Woo et al.’s dataset. A chromosome arm was defined as discordant if >=75% of the bins within that arm were discordant (**Methods**). Samples with <5% of the genome affected by CNAs were excluded from the analysis to avoid inflated discordance due to tumor impurity, and only weak correlation was observed between estimated tumor purity and genomic divergence in the remaining samples (**ED Fig. 1**).

**Figure 1:**
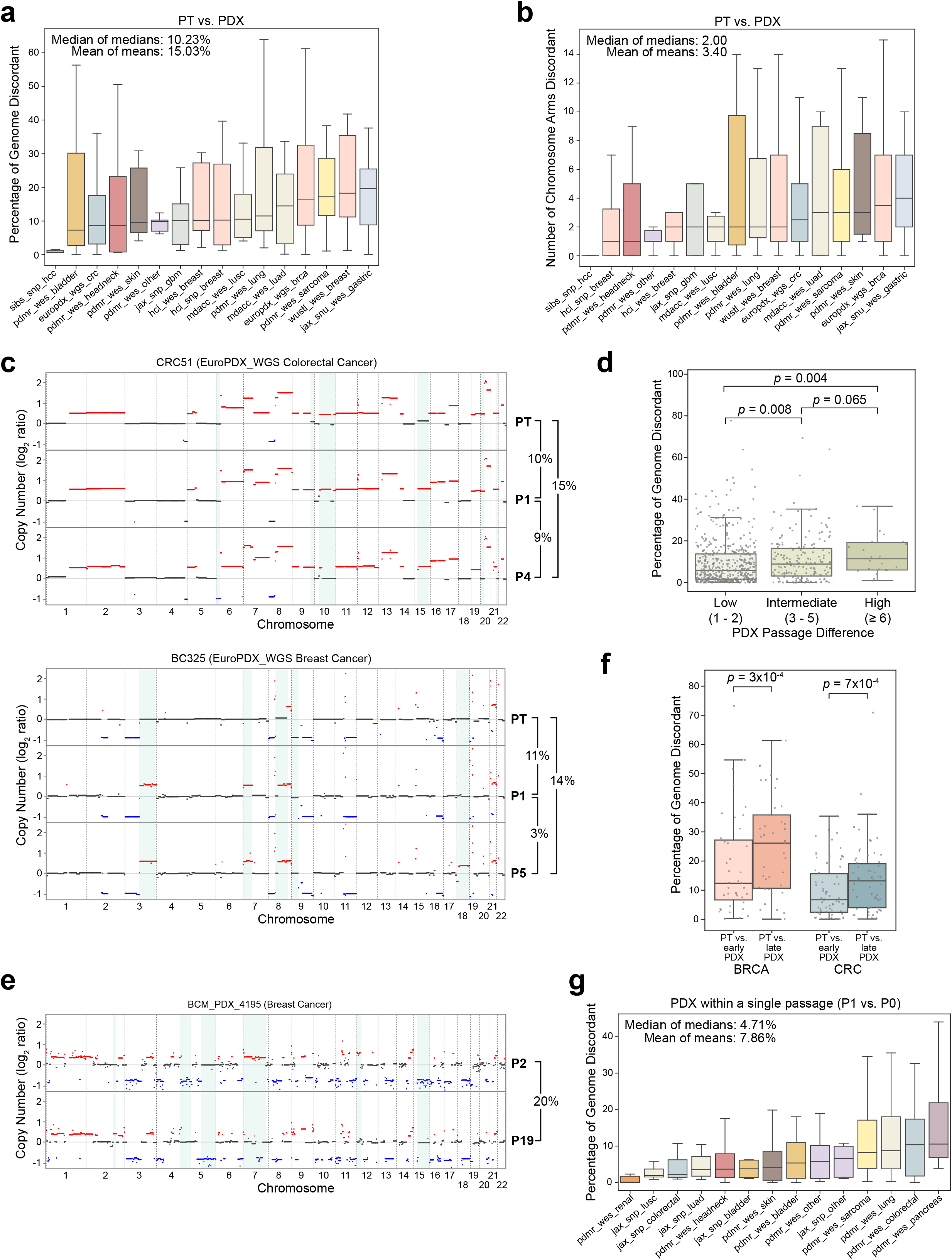
Thresholds-based comparison of the copy number landscapes of PTs and PDXs. (**a**) A comparison of the percent of the genome that is discordant between matched PT-PDX samples. In the median cohort, a median of 10.23% of the genome is altered between PTs and PDXs. Outliers were excluded from the plot. (**b**) A comparison of the number of chromosome arms that are discordant between matched PT-PDX samples. A median of 2 chromosome arms are altered between PTs and PDXs across cohorts. Outliers were excluded from the plot. (**c**) Representative examples of CN differences between matched PT, earlier-passage (P1) PDX, and later-passage (P4, P5) PDX samples from the EuroPDX_WGS colorectal and breast cancer cohorts. Red, CN gain; blue, CN loss. Prominent differences are highlighted with a light blue background. The fraction of the genome that is altered between samples is shown to the right of the plot. (**d**) A comparison of the percent of the genome that is discordant between matched samples of PDXs with a low (1-2), intermediate (3-5) or high (>=6) passage difference between them. The discordance increases with passage difference. P-values indicate significance from a Mann-Whitney U test. Circles, individual pairs. (**e**) A representative example of the CN differences between matched early passage (P2) PDX and late passage (P19) PDX samples from the BCM breast cancer cohort. Red, CN gain; blue, CN loss. Prominent differences are highlighted with a light blue background. The fraction of the genome that is altered between samples is shown to the right of the plot. (**f**) A comparison of the percent of the genome that is discordant between PTs vs. earlier-passage PDXs and PTs vs. later-passage PDXs, in breast and colorectal cancer cohorts that included matched ‘trios’ of PT and PDXs from two passages. P-values indicate significance from a one-sided Wilcoxon signed-rank test. Circles, individual pairs. (**g**) A comparison of the percent of the genome that is discordant between matched samples of PDXs at passage 0 and passage 1. A median of 4.71% of the genome is altered between P0 and P1 across cohorts. Bar, median; colored rectangle, 25th to 75th percentile; whiskers, Q1 – 1.5*IQR to Q3 + 1.5*IQR; outliers were excluded from the plot.

In line with our original analysis, we found that a median of 10.23% (range, 1.17% to 19.65%; mean of 15.03%) of the genome was differentially altered between PTs and PDXs across cohorts (**Fig. 1a** and **Supplementary Table 1**). Considering all PDX models from all cohorts together, a median of 10.95% (range 0% to 73.26%; mean of 15.56%) was differentially altered. Overall, over 25% of the genome was differentially affected by CN alterations in about a quarter (24%) of the matched PT-PDX samples (**Supplementary Table 1**). Many of the discrepancies were due to chromosome-arm aberrations that were present in only one of the paired samples: a median of 2 arm-level CN changes was differentially called between PTs and PDXs across cohorts (range, 0 to 4; mean of 3.40; **Fig. 1b**). The ichorCNA results showed similar discordance values with a median of 7.71% discordance (range 0.33% to 51.48%; mean of 15.41%; **ED Fig. 2a** and **Supplementary Table 2**) and 2 arm-level CN changes (range 0 to 10; mean 3.42; **ED Fig. 2b**). Therefore, although high overall correlations were observed in CN profiles between PTs and PDXs, such correlation values mask considerable differences between matched samples. Representative examples are shown in **Fig. 1c** and **ED Fig. 2c**. These findings demonstrate that the different magnitudes of PT-PDX discordance reported by our original study and by Woo et al. are not due to the different platforms that were used for CN calling, but rather due to our different definitions of ‘concordance’ and ‘discordance’.

A key question is whether genomic evolution with successive passaging leads to PDX diversification away from the genomic structure of primary tumors. Woo et al. did not identify a downward concordance trend over passaging, concluding that late PDX passages maintain CN profiles similar to early PDX passages^2^. However, using our measures for CN discordance (rather than the Pearson’s correlation used by Woo et al.), we find that the CN discordance between matched pairs of PDXs significantly increases with passaging. In other words, the higher the passage difference, the more discordant matched samples are **(Fig. 1d** and **ED Fig. 2d**). Notably, the BCM breast cancer cohort is composed of pairs of PDXs 17 to 21 passages apart, the highest passage differences in the entire Woo et al. dataset; by our analysis, this cohort also has the highest median discordance between paired PDXs of any cohort (median, 10.79%; range, 5.61% to 36.60%; mean, 13.3%), and much of this difference is due to aneuploidy (i.e., chromosome-arm or whole-chromosome alterations; **Fig. 1e**, **ED Fig. 3a** and **Supplementary Tables 1 and 2**). Moreover, in cohorts that include ‘trios’ of PTs and a PDX model evaluated at two different passages, the discordance between PTs and later-passage PDXs was significantly higher than that between PTs and earlier-passage PDXs (p=0.0003 and p=0.0007, one-sided Wilcoxon signed-rank test; for the breast and colon cohorts, respectively; **Fig. 1f** and **ED Fig. 3b**). Additionally, across all cohorts, a median of 4.71% (range, 0.093% to 10.50%; mean, 7.86%) of the genome was altered between passage #0 and passage #1 of the PDX models (**Fig. 1g** and **ED Fig. 4a-c)**, consistent with our previous observation that much of the genomic evolution occurs within the first few PDX passages. These findings provide strong evidence that genomic evolution leads to gradual PDX diversification.

Woo et al. performed a GISTIC analysis to identify recurrent CNAs in primary tumors and in PDXs. This analysis did not identify recurrent alterations of a specific locus, gene or pathway during PDX engraftment or passaging, and the authors interpreted this as evidence for lack of selection. In our previous study, we also did not identify specific CNAs that were selected for in the PDX models^1^. This is not surprising, as both studies are limited by the number of samples when controlling for tumor type. We confirmed that the sample sizes in the study by Woo et al. are not powered (power < 80%) to detect a 10% prevalence decrease for 96% of the TCGA recurrent aneuploidies (**ED Fig. 5**). Therefore, the inability of both papers to identify specific recurrent mouse-induced CNAs should not be considered as a proof for the absence of selection, and much larger cohorts will be necessary to further address this question.

Importantly, however, clonal selection does not necessarily have to involve recurrent events. We previously found that when the same PT is transplanted into different mice, the ‘sibling’ PDX models tend to acquire the same CNAs, indicating positive selection^1^. Moreover, we previously reported that the rate of CN evolution in PDXs is strongly correlated with the intra-tumor heterogeneity of the primary tumor types. This result is mirrored by Woo et al.’s observation that the CN differences between PTs and PDXs are similar to the differences across multi-region tumor samples. We note that the observation that PDX genomic evolution is largely driven by spatial heterogeneity by no means diminishes its potential consequences; PDX evolution due to spatial heterogeneity could still lead to considerable differences between PTs and their derived PDXs. While we agree that “the goal of a successful treatment would be to eradicate all of the multiple regions of the tumor”, considerable PT-PDX differences in the clonal composition of the tumor may reduce the utility of the model. Therefore, the finding that the differences between PTs and PDXs are no greater than those observed spatially within patients’ solid tumors does not alleviate the concerns about the genetic stability of the PDX system, especially in the context of ‘tumor avatars’.

In summary, our analysis of the dataset presented by Woo et al. constitutes a DNA-based validation of our previous results. Importantly, the disagreement between the studies is not a matter of the method used for CN calling, but rather a question of data analysis and interpretation. We believe that the controversy can be boiled down to the following questions: Is it alarming or negligible that a median of 10% of the genome is different between PTs and PDXs, that these CN alterations often include chromosome-arm gains and losses, and that the genomic similarity of PDXs to their PT of origin decreases over time? And, perhaps the most important question: How do such changes affect the phenotypic stability of the models, and their ability to serve as accurate ‘tumor avatars’? Addressing these questions directly and systematically is important for optimizing the use of these unique and valuable cancer models. Until then, we strongly support Woo et al.’s final recommendation “to confirm the existence of expected molecular targets and obtain sequence characterizations in the cohorts used for [drug] testing as close to the time of treatment study as is practical”.

## Online Methods

### PDX dataset and preprocessing

Segments of log_2_(CN ratio) values across the genome for both PT and PDX samples were acquired from Woo, et al^2^. Samples whose CN profiles had been estimated from SNP arrays, whole-exome sequencing data, and low-pass whole-genome sequencing data were included, while samples whose profiles had been estimated from RNA-sequencing and gene expression microarray data were excluded, to avoid potential issues with expression-based CN inference. The final dataset consisted of 1,429 samples across 33 cohorts containing at least one pair of matched samples.

The log_2_(CN ratio) segments for each sample were binned into 1 Mb windows across the genome. Bins that overlapped more than one segment were assigned the value of the overlapping segment with greatest absolute value.

### Sample pairing scheme

#### PT vs. PDX and PDX vs. PDX discordance across different cohorts

Each PT sample was compared to each of its available PDXs. Each PDX sample was compared to every other PDX available from the same original PT, provided that the PDX samples were of different passage numbers. Cohorts with <5 PT-PDX or PDX-PDX pairs were excluded from the respective analyses.

#### PDX vs. PDX discordance by passage difference

PDX samples were paired with later-passage PDX samples in the same PDX model with the minimum possible passage difference.

#### Trios analysis of EuroPDX WGS cohorts

For each model with a PT and two PDX samples of different passage numbers, the PT was compared to both PDXs to calculate the discordance between PT and the earlier-passage PDX and that between the PT and the later-passage PDX.

### Computing CNA discordance between paired samples using thresholds

CN gains and losses were defined as log_2_(CN ratio) >= 0.3 and log_2_(CN ratio) <= −0.3, respectively. For a 1 Mb bin to be discordant between two samples, one sample must uniquely have a CN gain or loss, and additional criteria specifying the difference in log_2_(CN ratio) required between the two samples must be satisfied, to exclude borderline cases. In full, if two samples have log_2_(CN ratio) values A and B, the samples are discordant if at least one of the following conditions are met:

- (A >= 0.3 & B < 0.3) & ((A – B) >= 0.3 | (B < 0.1))
- (A <= −0.3 & B > −0.3) & ((A – B) <= −0.3 | (B > −0.1))
- (B >= 0.3 & A < 0.3) & ((B – A) >= 0.3 | (A < 0.1))
- (B <= −0.3 & A > −0.3) & ((B – A) <= −0.3 | (A > −0.1))

The fraction of 1 Mb bins discordant across the genome between two samples is defined as the number of discordant bins divided by the number of bins across the genome for which both samples have log_2_(CN ratio) values.

For a chromosome arm to be discordant between two samples, >= 75% of the 1 Mb bins in the chromosome arm must be discordant using the above thresholds, and these discordances must always be due to one sample having the greater log_2_(CN ratio) and the other sample having the lesser log_2_(CN ratio).

To exclude highly-discordant cases that may be due to tumor purity issues, a sample was assumed to lack sufficient tumor fraction to analyze if < 5% of the 1 Mb bins in its genome with log_2_(CN ratio) values had absolute values of >= 0.3. 100 samples were removed from analysis due to this criterion.

### Copy number analysis using ichorCNA

For each sample, the log_2_(CN ratio) values at each 1 Mb bin were passed in as input to a modified version of the Hidden Markov Model-based algorithm ichorCNA(11). This version of ichorCNA calls large-scale CNAs and estimates tumor fraction and ploidy from a sample’s median-normalized, GC-corrected log_2_(CN ratio) values across the genome. ichorCNA uses a probabilistic model to predict CNAs and does not use log_2_(CN ratio) thresholds. PT samples were run with normal cell fraction initializations of 0.25, 0.5, and 0.75, and PDX samples were initialized with 0.05 normal cell fraction. Samples were run for starting conditions of both ploidy 2 and ploidy 3. ichorCNA selects a sample’s final solution based on the likelihood scores of the set of solutions produced from the different runs.

If the ploidies of each set of paired samples in a model did not converge to within 0.2 of each other, some of the samples were rerun with their starting ploidies restricted to one value to increase the likelihood of each sample having similar estimated ploidy. The details of this approach depended on the composition of the model: If a model was composed of one PT and one PDX, the PT sample was rerun with the closest possible starting ploidy to the PDX’s estimated ploidy. If a model was composed of two PDXs, the sample with lower tumor fraction was rerun with the closest possible starting ploidy to its pair’s estimated ploidy, and if these samples had equal tumor fraction, the sample with higher ploidy was rerun. Finally, if a model was composed of more than two samples, the starting ploidy was restricted to the most common rounded estimated ploidy. In the case of multiple equally common rounded estimated ploidies, the new starting ploidy was picked randomly from those ploidies.

### Computing CNA discordance between paired samples from ichorCNA results

CNA profiles across the genome were transformed to ploidy 2 space using the equation:

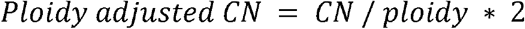

Rounded, ploidy-adjusted CN of > 2 was defined as a CN gain, and < 2 as a loss. For a 1 Mb bin to be discordant between two samples, one sample must uniquely have a CN gain or loss, and the unrounded ploidy-adjusted CNs must also differ between the two samples by >= 0.5.

ichorCNA calls some 1 Mb bins as subclonal CN gains/losses. Subclonal CN calls with > 90% estimated cellular prevalence were treated as clonal gains/losses, and subclonal CN calls with < 10% cellular prevalence were treated as neutral CN. These calls then underwent the same ploidy-adjustment as other clonal calls.

Subclonal CN calls with intermediate cellular prevalence were not considered to be discordant from either neutral copy number or from the gain/loss they had character of.

Chromosome arm level discordance between two samples was again defined as >= 75% of the 1 Mb bins in an arm with data being discordant due to one sample always having the greater CN and the other sample having the lesser CN.

Samples were excluded from analysis if ichorCNA estimated the sample contained < 5% tumor fraction and/or if < 5% of the bins were clonally altered from neutral CN after ploidy-adjustment and rounding. 52 samples were excluded based on these criteria.

### Statistical analysis

The statistical significance of the difference in genomic discordance between PTs vs. earlier-passage PDXs and PTs vs. later-passage PDXs was determined by one-sided Wilcoxon signed-rank test. The statistical significance of the difference in genomic discordance between matched PDX pairs with low (1-2), intermediate (3-5) or high (≥6) passage number between them was determined by Mann-Whitney U test. Statistical analyses were performed using the SciPy Python library. Plotting was performed using the Pandas and Seaborn Python libraries.

### Power analysis for PDX selection against recurrent aneuploidies analysis

The prevalence of specific aneuploidies in patient tumors for a given tumor type was calculated from the aneuploidy calls of 10,522 cancer genomes from The Cancer Genome Atlas (TCGA) as presented in Taylor et al.^12^. Arm-level alterations with prevalence > 0.25 were defined as recurrent. We determined the minimum sample size n needed to detect a 10% absolute decrease in the prevalence of a recurrent arm for a specific tumor type with sufficient power of 80%, using a one-sided one-sample proportion test.

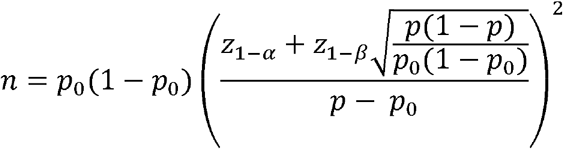

The reference p_0_ represents the expected prevalence of the recurrent arm in the TCGA patient tumors for a specific tumor type, while p represents the scenario for a 10% prevalence decrease, p = p_0_ −0.1. z is the inverse of the cumulative distribution function with Type I error α = 0.05, Type II error β = 0.2, and power 1-β=0.8. For each recurrent arm of a given tumor type, we compared the powered sample size n to the number of PT-PDX comparisons available in our dataset for the given tumor type.

## Code Availability

The code used to analyze the data are publicly available, or available upon request.

## Data availability

All datasets used in this analysis are publicly available.

## Acknowledgments

U.B.-D. is supported by the Azrieli Foundation (U.B.-D.), the Richard Eimert Research Fund on Solid Tumors (U.B.-D.), the Tel-Aviv University Cancer Biology Research Center (U.B.-D.), the Israel Cancer Association (U.B.-D.), the DoD CDMRP career development award (grant #CA191148), the BSF project grant (#2019228), and the ICRF Gesher Award (U.B.-D.). G.H is supported by the NIH/NCI Transition Career Development Award (K22 CA237746) and The V Foundation V Scholar Grant. Research in the Ha lab is funded in part through the NIH/NCI Cancer Center Support Grant (P30 CA015704) and ORIP Grant (S10OD028685).

## Author Contributions

U.B.-D. and G.H. conceived the analysis. A.H., M.G., and G.H. performed the analyses and generated the plots and the tables. All authors participated in analyzing the data and writing the manuscript. U.B.-D and G.H. supervised the project.

## Figure Legends

**Extended data Figure 1: The contribution of purity to CN discordance.** (**a**) The Spearman correlation between the PT purity estimate and the percent of the genome that is discordant between the PT and its matched PDX in the thresholds-based CN analysis. (**b**) The Spearman correlation between the PT purity estimate and the percent of the genome that is discordant between the PT and its matched PDX in the ichorCNA-based CN analysis. Data points correspond to PT-PDX matched samples.

**Extended Data Figure 2: ichorCNA-based comparison of the copy number landscapes of PTs and PDXs.** (**a**) A comparison of the percent of the genome that is discordant between matched PT-PDX samples. In the median cohort, a median of 7.71% of the genome is altered between PTs and PDXs. Outliers were excluded from the plot. (**b**) A comparison of the number of chromosome arms that are discordant between matched PT-PDX samples. In the median cohort, a median of 2 chromosome arms are altered between PTs and PDXs. Outliers were excluded from the plot. (**c**) Representative examples of ichorCNA results showing the CN differences between matched PT, earlier-passage (P1) PDX and later-passage (P5) PDX samples from the EuroPDX_WGS colorectal and breast cancer cohorts. Red, CN gain; blue, CN loss. Prominent differences are highlighted with a light blue background. The fraction of the genome that is altered between samples is shown to the right of the plot. (**d**) A comparison of the percent of the genome that is discordant between matched samples of PDXs with a low (1-2), intermediate (3-5) or high (≥6) passage difference between them. The discordance increases with larger passage differences. P-values indicate significance from a Mann-Whitney U test. Circles, individual pairs. Bar, median; colored rectangle, 25th to 75th percentile; whiskers, Q1 – 1.5*IQR to Q3 + 1.5*IQR.

**Extended Data Figure 3: ichorCNA-based comparison of the copy number landscapes of PTs and PDXs (continued).** (**a**) A representative example of the CN differences between matched early passage (P1) PDX and late passage (P19) PDX samples from the BCM breast cancer cohort. Red, CN gain; blue, CN loss. Prominent differences are highlighted with a light blue background. The fraction of the genome that is altered between samples is shown to the right of the plot. (**b**) A comparison of the percent of the genome that is discordant between PTs vs. earlier-passage PDXs and PTs vs. later-passage PDXs, in breast and colorectal cancer cohorts that included matched ‘trios’ of PT and PDXs from two passages. P-values indicate significance from a one-sided Wilcoxon signed-rank test. Circles, individual pairs; bar, median; colored rectangle, 25th to 75th percentile; whiskers, Q1 – 1.5*IQR to Q3 + 1.5*IQR.

**Extended Data Figure 4: Comparison of the fraction of the genome that is altered within a single passage.** (**a**) A comparison of the percent of the genome that is discordant between matched samples of PDXs at passage 0 and passage 1 with the ichorCNA-based analysis. In the median cohort, a median of 3.85% of the genome is altered between P0 and P1. Bar, median; colored rectangle, 25th to 75th percentile; whiskers, Q1 - 1.5*IQR to Q3 + 1.5*IQR; outliers were excluded from the plot. (**b**) The distribution of the passage difference between matched PDX samples across the Woo et al. cohorts, which were used for the thresholds-based analysis. (**c**) The distribution of the passage difference between matched PDX samples across the Woo et al. cohorts, which were used for the ichorCNA-based analysis.

**Extended Data Figure 5: Power analysis to determine the required sample size to detect selection against recurrently altered chromosome arms.** Chromosome arm aneuploidy prevalence of TCGA tumors (12) are shown as points for 13 tumor types (x-axis); recurrent armlevel aneuploidies with prevalence >= 0.25 are shown. The observed sample sizes of the corresponding tumor types in the Woo et al. study are indicated by the y-axis. Points above the black curve (green shading) are statistically powered to detect a 10% decrease in absolute prevalence from PTs to PDXs (with power of 80% and α=0.05), while points below the curve (red shading) are not. Statistical power analysis was performed using a single-sample, one-sided proportion test (**Methods**). Select recurrently altered chromosome arms are labeled.

## Table Legends

**Supplementary Table 1: Thresholds-based CN discordance values.** Shown are all pairs that met the quality control requirements (**Methods**) and were used for the thresholds-based discordance analyses.

**Supplementary Table 2: ichorCNA-based CN discordance values.** Shown are all pairs that met the quality control requirements (Methods) and were used for the ichorCNA-based discordance analyses.

## References

1. Ben-David, U. et al. Patient-derived xenografts undergo mouse-specific tumor evolution. Nat. Genet. (2017) doi:10.1038/ng.3967.

2. Woo, X. Y. et al. Conservation of copy number profiles during engraftment and passaging of patient-derived cancer xenografts. Nat. Genet. 53, 86–99 (2021).

3. Davies, N. J. et al. Dynamic changes in clonal cytogenetic architecture during progression of chronic lymphocytic leukemia in patients and patient-derived murine xenografts. Oncotarget (2017) doi:10.18632/oncotarget.17432.

4. Gendoo, D. M. A. et al. Whole genomes define concordance of matched primary, xenograft, and organoid models of pancreas cancer. PLoS Comput. Biol. (2019) doi:10.1371/journal.pcbi.1006596.

5. Van Der Heijden, M. et al. Spatiotemporal regulation of clonogenicity in colorectal cancer xenografts. Proc. Natl. Acad. Sci. U. S. A. (2019) doi:10.1073/pnas.1813417116.

6. Liu, Y. et al. Gene expression differences between matched pairs of ovarian cancer patient tumors and patient-derived xenografts. Sci. Rep. (2019) doi:10.1038/s41598-019-42680-2.

7. Tew, B. Y. et al. Patient-derived xenografts of central nervous system metastasis reveal expansion of aggressive minor clones. Neuro. Oncol. (2020) doi:10.1093/neuonc/noz137.

8. Vaubel, R. A. et al. Genomic and phenotypic characterization of a broad panel of patient-derived xenografts reflects the diversity of glioblastoma. Clin. Cancer Res. (2020) doi:10.1158/1078-0432.CCR-19-0909.

9. Kim, J., Rhee, H., Kim, J. & Lee, S. Validity of patient-derived xenograft mouse models for lung cancer based on exome sequencing data. Genomics and Informatics (2020) doi:10.5808/GI.2020.18.1.e3.

10. Peng, D. et al. Evaluating the transcriptional fidelity of cancer models. bioRxiv 2020.03.27.012757 (2021) doi:10.1101/2020.03.27.012757.

11. Adalsteinsson, V. A. et al. Scalable whole-exome sequencing of cell-free DNA reveals high concordance with metastatic tumors. Nat. Commun. (2017) doi:10.1038/s41467-017-00965-y.

12. Taylor, A. M. et al. Genomic and Functional Approaches to Understanding Cancer Aneuploidy. Cancer Cell 33, 676–689 e3 (2018).

