## Supplementary figures and images for "DNA-based copy number analysis confirms genomic evolution of PDX models"

### Extended Data Figure 1

**a**

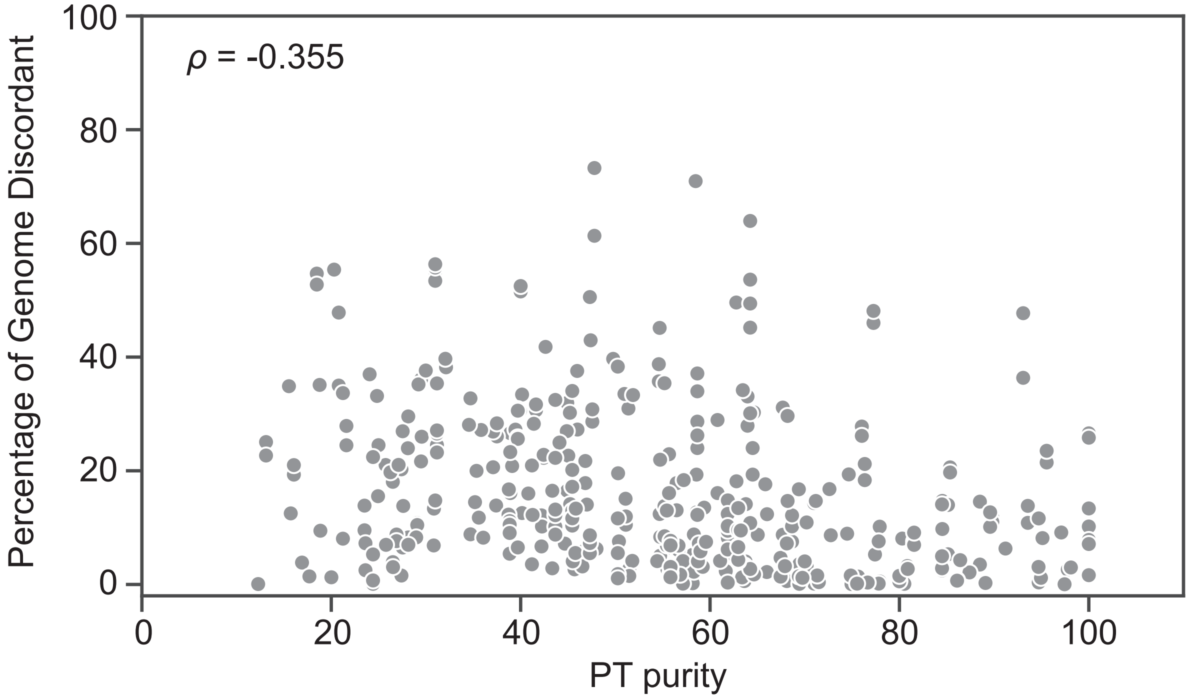


**b**

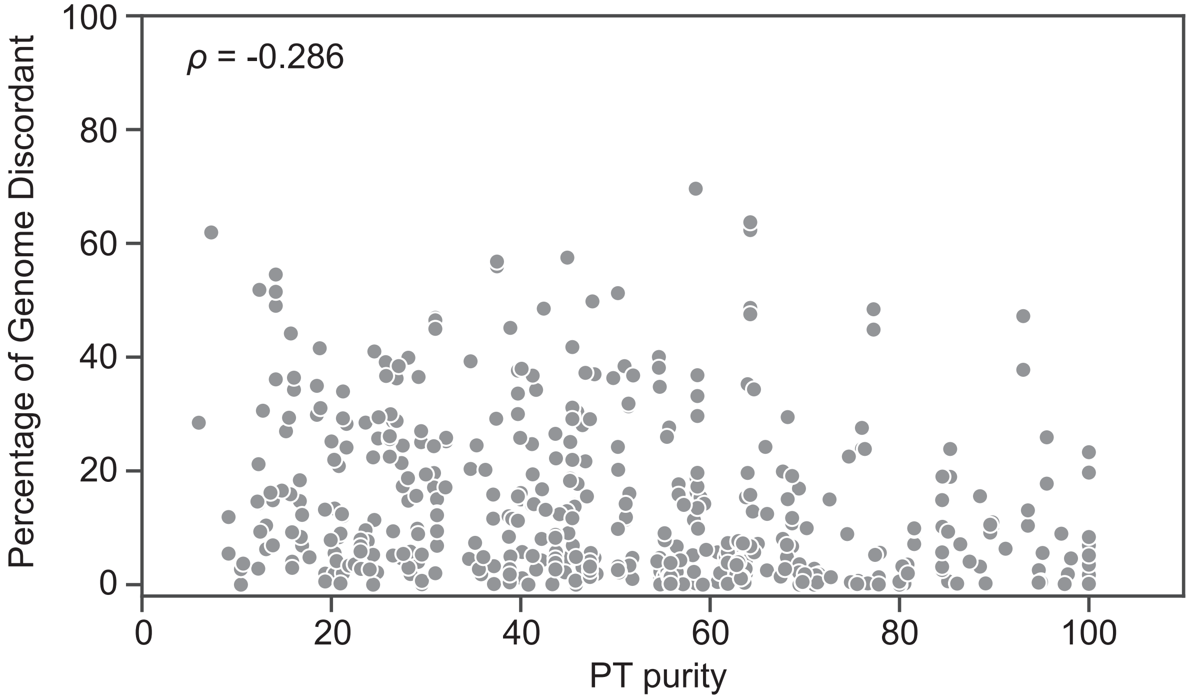


**Extended Data Figure 1**

### Extended Data Figure 2

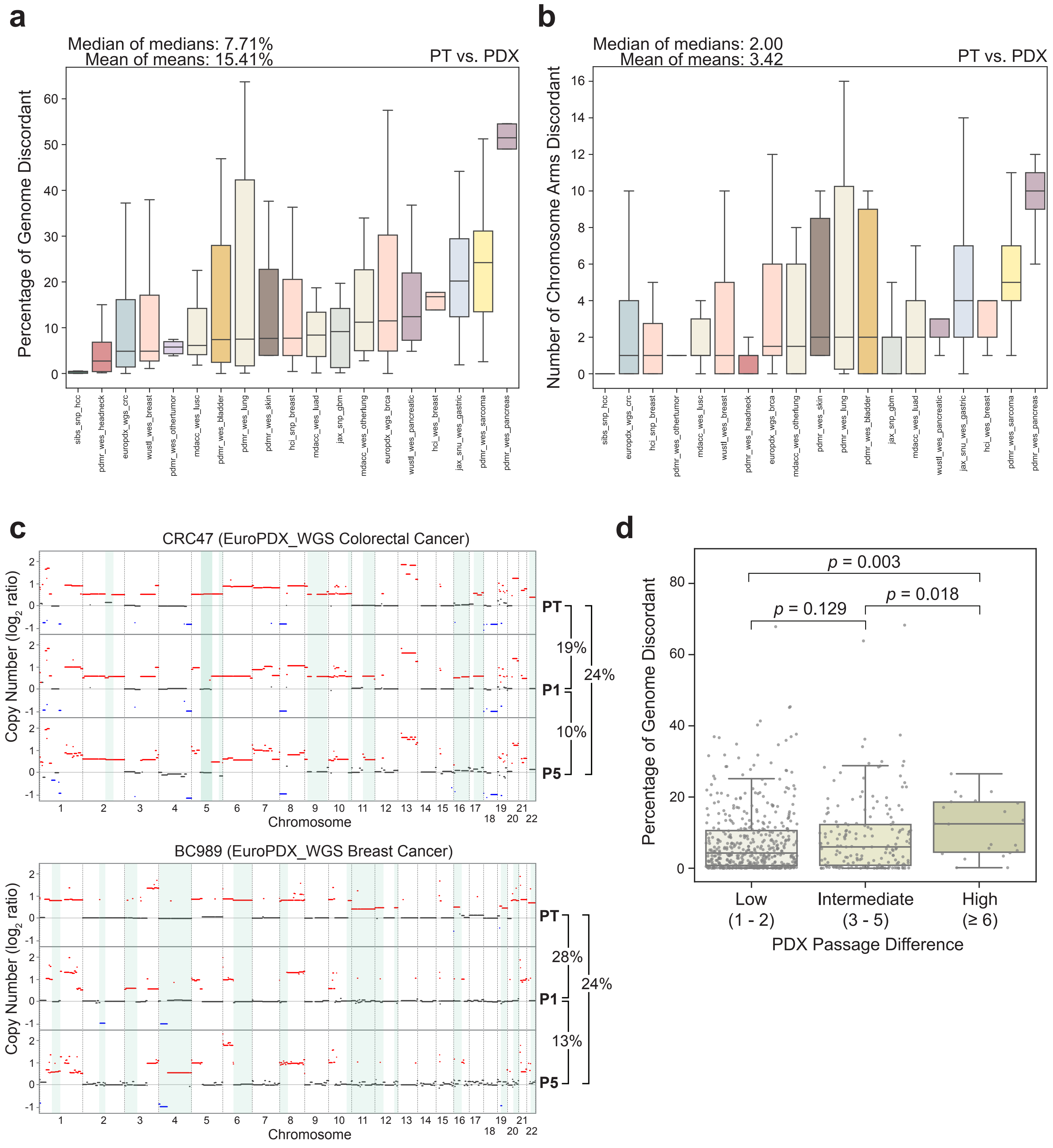


#### Extended Data Figure 2

### Extended Data Figure 3

**a**

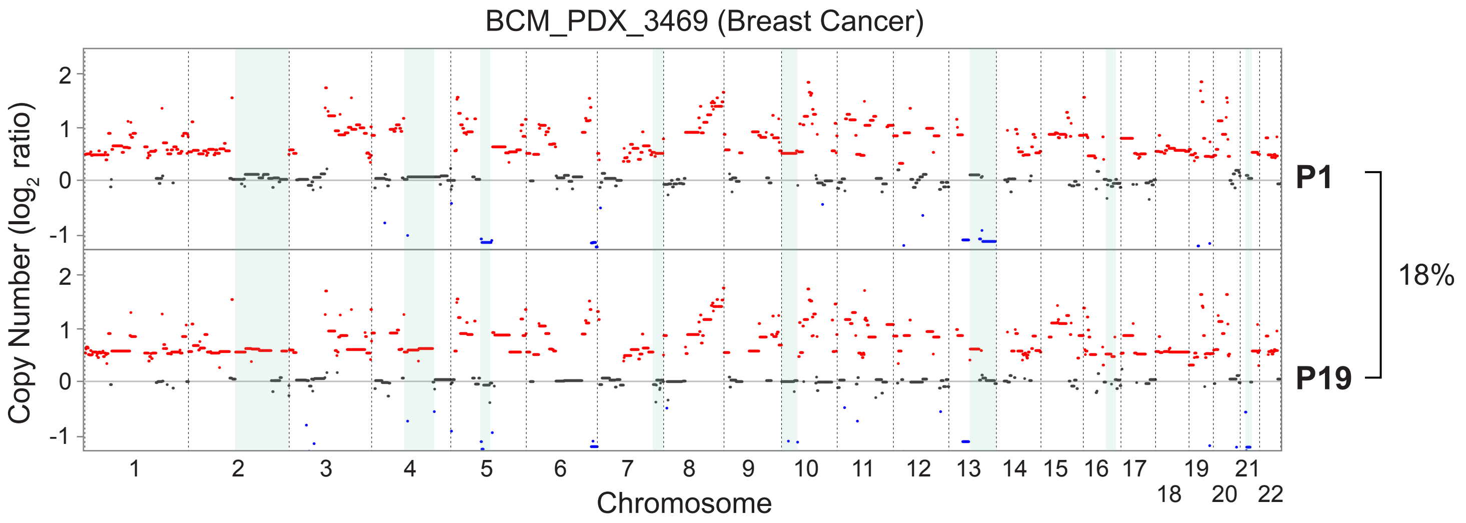


**b**

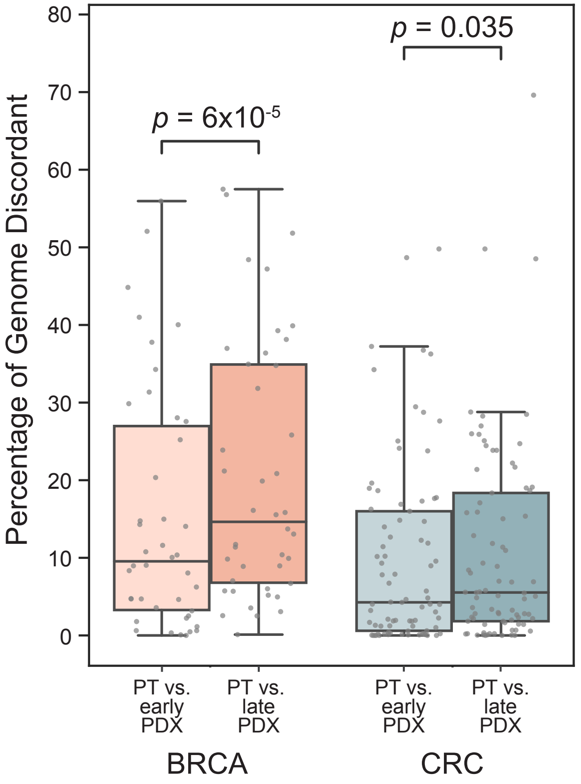


**Extended Data Figure 3**

### Extended Data Figure 4

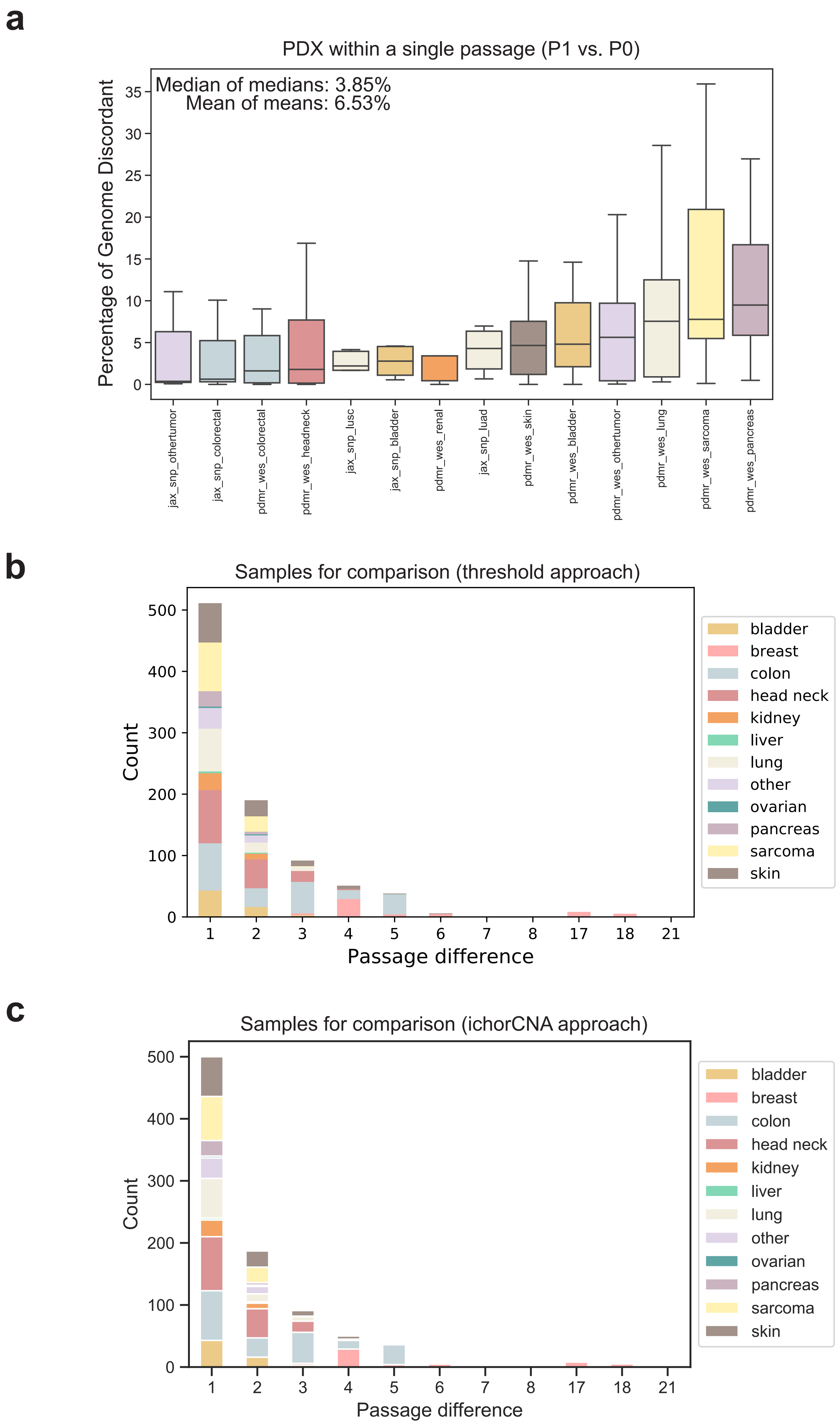


**Extended Data Figure 4**

### Extended Data Figure 5

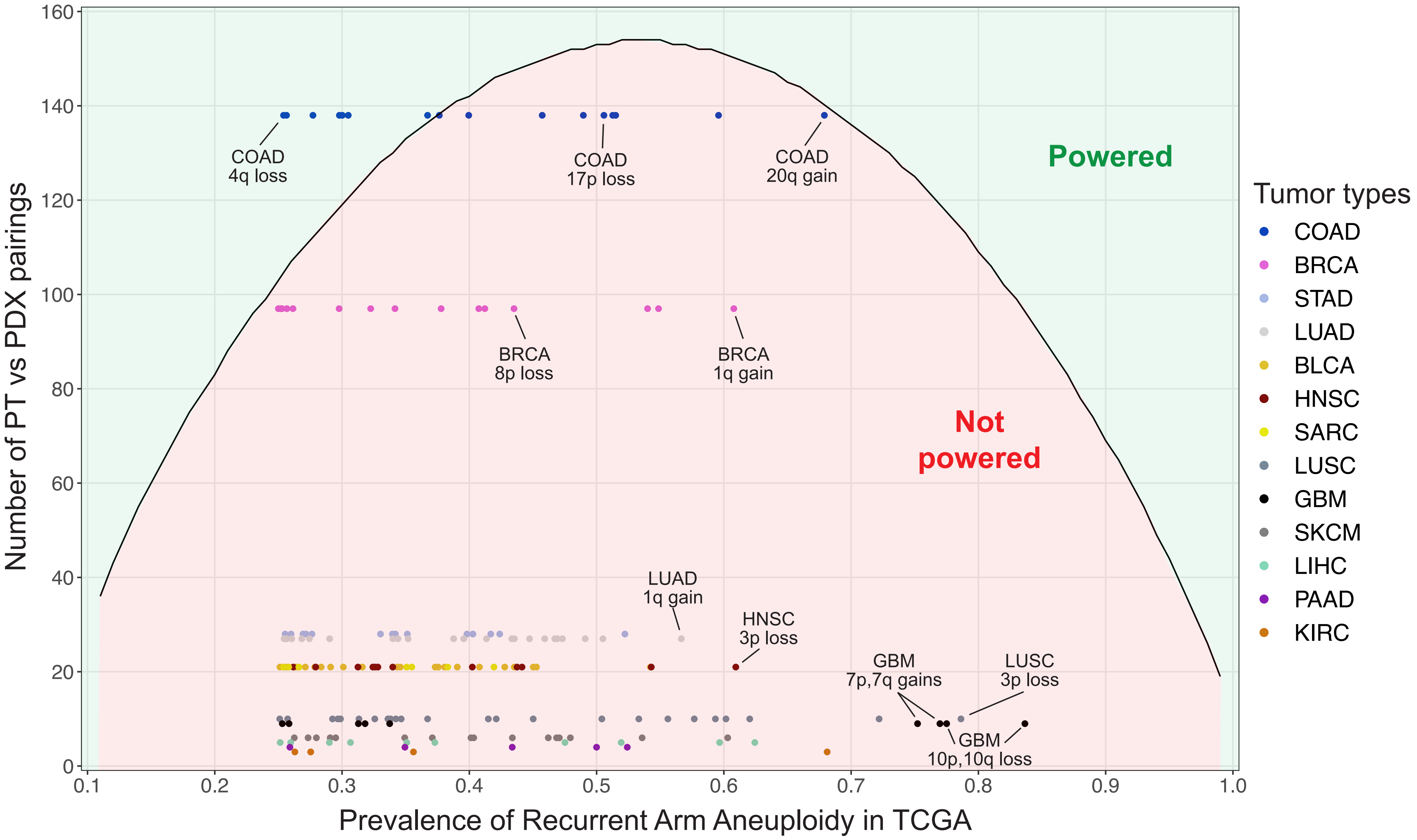


**Extended Data Figure 5**
